# FCHSD2 Controls Oncogenic ERK1/2 Signaling Outcome by Regulating Endocytic Trafficking

**DOI:** 10.1101/2020.01.29.924449

**Authors:** Guan-Yu Xiao, Sandra L. Schmid

## Abstract

Cancer progression is driven, in part, by altered signaling downstream of receptor tyrosine kinases (RTKs). Surface expression and RTK activity are regulated by clathrin-mediated endocytosis (CME), endosomal recycling or degradation. In turn, oncogenic signaling downstream of RTKs can reciprocally regulate endocytic trafficking, creating feedback loops that enhance tumor progression. We previously reported a cancer-cell specific function of FCHSD2 (FCH/F-BAR and double SH3 domain-containing protein) in regulating CME in non-small-cell lung cancer (NSCLC) cells. Here, we report that FCHSD2 loss impacts recycling of EGFR and MET, diverting their trafficking toward late endosomes and lysosomes. FCHSD2 depletion results in the nuclear translocation of active ERK1/2, leading to enhanced transcription and upregulation of EGFR and MET. The small GTPase, Rab7, is essential for the FCHSD2 depletion-induced effects. Correspondingly, FCHSD2 loss correlates with higher tumor grades of NSCLC. Clinically, NSCLC patients expressing high FCHSD2 exhibit elevated survival, whereas patients with high Rab7 expression display decreased survival rates. Our study provides new insight into the molecular nexus for crosstalk between oncogenic signaling and RTK trafficking that controls cancer progression.

## Introduction

Non-small-cell lung cancer (NSCLC) is the major world-wide cause of death from cancer (Siegel, Miller et al., 2019). Transformed NSCLC cells surreptitiously multiply while undergoing generations of selected evolution to acquire the characteristics of an aggressive and metastatic tumor, in part driven by altered signaling, which is associated with activation of receptor tyrosine kinases (RTKs) (Bacac & Stamenkovic, 2008, Gower, Wang et al., 2014, Hanahan & Weinberg, 2011). The expression and activity of cell surface RTKs, in turn, is regulated predominantly through clathrin-mediated endocytosis (CME) (Conner & Schmid, 2003, Gonnord, Blouin et al., 2012, McMahon & Boucrot, 2011), endosomal recycling or degradation (Mellman & Yarden, 2013, Paul, Jacquemet et al., 2015, Sigismund, Confalonieri et al., 2012). Hence, a link between endocytic trafficking and cancer progression has been suggested (Lanzetti & Di Fiore, 2008, Mellman & Yarden, 2013, Mosesson, Mills et al., 2008). Yet, few studies have focused on cancer cell specific alterations in endocytic trafficking.

CME is the major endocytic pathway that determines the rates of internalization of plasma membrane receptors, regulates their expression on the cell surface, and controls their downstream signaling activities (Conner & Schmid, 2003, Gonnord et al., 2012, McMahon & Boucrot, 2011). We previously discovered that oncogenic signaling downstream of surface RTKs can, in turn regulate CME and early recycling pathways by creating feedback loops that influence signaling, migration and metastasis in NSCLC cells (Chen, Bendris et al., 2017, Schmid, 2017, Xiao, Mohanakrishnan et al., 2018). Moreover, some of these mechanisms for reciprocal crosstalk between signaling and the endocytic trafficking pathway appear to be specific for, or co-opted by cancer cells to enhance tumor progression (Schmid, 2017, Xiao et al., 2018). We have termed these cancer-specific changes in the endocytic machinery ‘adaptive’ endocytic trafficking and hypothesize that these ‘gain-of-function’ changes in endocytic trafficking contribute to cancer progression and metastasis (Schmid, 2017).

The mechanisms that control the crosstalk between cargo (especially signaling receptors) and the endocytic machinery and their roles in cancer have not been explored. We recently discovered that the cancer-specific activation of FCHSD2 (FCH/F-BAR and double SH3 domain-containing protein) downstream of ERK1/2 contributes to adaptive CME in NSCLC cells (Xiao et al., 2018), in this case by suppressing EGFR signaling. Similarly, its *Drosophila* ortholog, Nervous Wreck (*Nwk*) suppresses BMP signaling, but by regulating endosomal recycling at the synapse (Rodal, Blunk et al., 2011). Accordingly, it is now critical to determine whether FCHSD2 also functions in endosomal trafficking of RTKs to regulate their oncogenic signaling from endosomes and whether this effects human tumor progression.

To address this issue, we used HCC4017 and H1975 NSCLC cells, which exhibit oncogenic signaling pathways downstream of Kirsten Ras (KRas^G12C^) or EGFR^T790M/L858R^ mutations, respectively. Here, we measured the effects of FCHSD2 depletion on the endocytic recycling and trafficking of the RTKs, EGFR and MET, and the consequences of these alterations on downstream signals. We demonstrated that FCHSD2 functions as a switch to regulate the trafficking pathway and destination of the RTKs through negative regulation of the small GTPase, Rab7. FCHSD2-dependent RTK trafficking controls the nuclear translocation of ERK1/2 signaling and expression of the RTKs. Our study provides a novel mechanism of action by which protein traffic between endosomal compartments controls the outcome of ERK1/2 signaling and to affect NSCLC progression.

## Results

### FCHSD2 regulates endosomal trafficking of TfnR and EGFR in NSCLC cells

To test whether FCHSD2, like its *Drosophila* homologue also functions in endosomal trafficking, we first assessed recycling of transferrin receptor (TfnR), a canonical marker for the quantification of endosomal trafficking (Harding, Heuser et al., 1983). To further determine which step(s) are affected, we measured TfnR recycling directly from early endosomes, following a 10 min pulse of internalized ligand or through perinuclear recycling endosomes following a 30 min pulse (Maxfield & McGraw, 2004). FCHSD2 knockdown (KD) by siRNA selectively inhibits slower recycling of TfnR, presumably via recycling endosomes (Fig. 1A and B). However, unlike the cancer cell-specific role of FCHSD2 in regulating CME downstream of ERK1/2 (Xiao et al., 2018), its function in endosomal recycling was neither dependent on ERK1/2 activity nor cancer cell-specific (EV Fig. 1).

**Figure 1.**
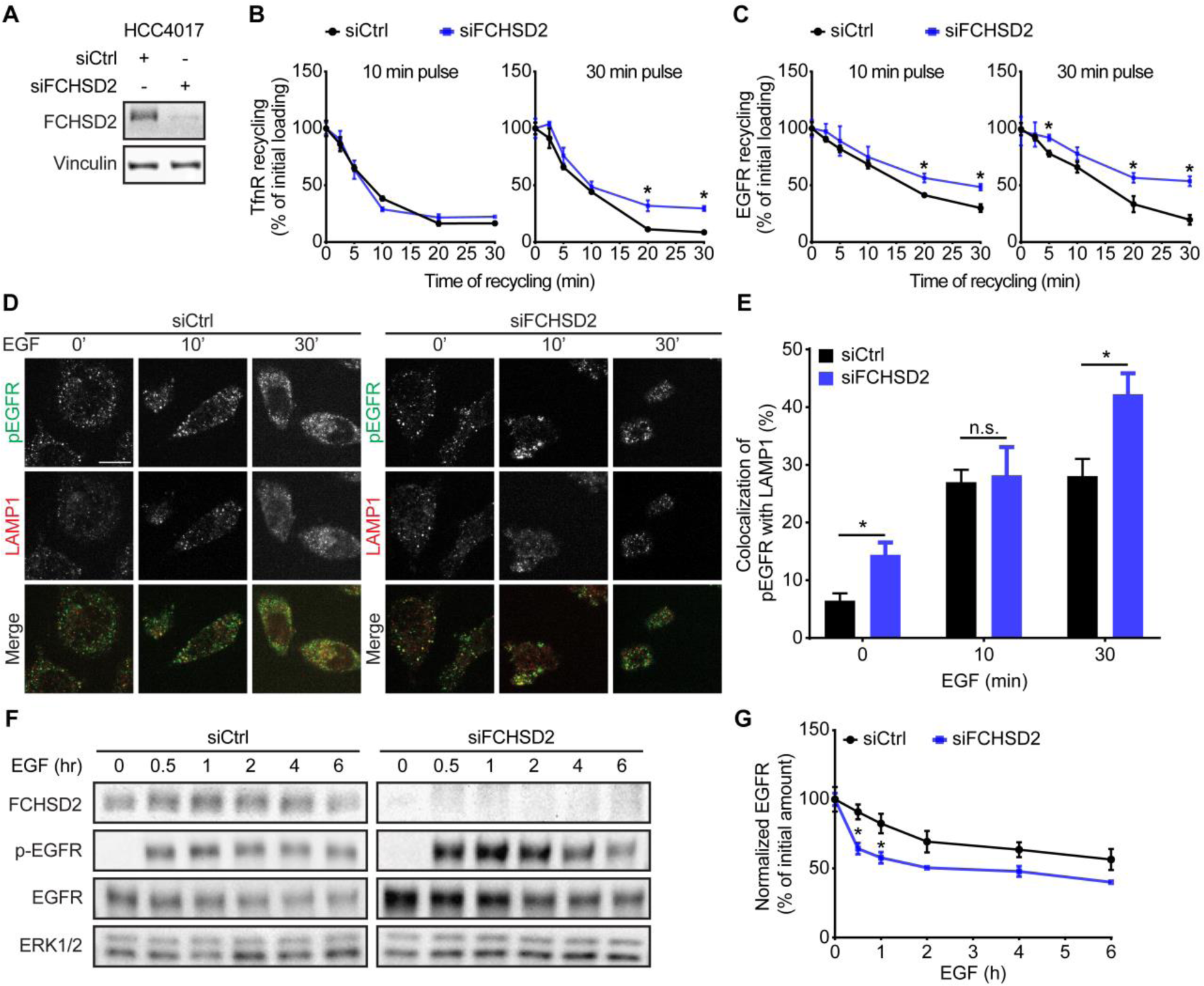
FCHSD2 regulates TfnR and EGFR endocytic trafficking in NSCLC cells. **A,** The knockdown of FCHSD2 in control or FCHSD2 siRNA-treated HCC4017 cells. **B** and **C,** Endocytic recycling of TfnR (**B**) or EGFR (**C**) was measured in control or FCHSD2 siRNA-treated HCC4017 cells. Cells were pulsed for 10 min or 30 min with 10 μg/ml biotinylated Tfn (**B**) or 20 ng/ml biotinylated EGF (**C**), stripped, and reincubated at 37°C for the indicated times before measuring the remaining intracellular Tfn or EGF. Percentage of recycled biotinylated Tfn or EGF was calculated relative to the initial loading. Data represent mean ± SEM (*n* = 3). Two-tailed Student’s *t* tests were used to assess statistical significance versus siCtrl. \**P* < 0.05, \*\**P* < 0.005, \*\*\**P* < 0.0005. **D,** Representative confocal images of pEGFR and LAMP1 immunofluorescence staining in control or FCHSD2 siRNA-treated HCC4017 cells. Cells were incubated with 20 ng/ml EGF for 30 min at 4°C, washed, and reincubated at 37°C for the indicated times. Scale bar, 12.5 μm. **E,** Colocalization of pEGFR and LAMP1 immunofluorescence staining in the cells as described in **D**. Data were obtained from at least 40 cells in total/condition and represent mean ± SEM. Two-tailed Student’s *t* tests were used to assess statistical significance. n.s., not significant, \**P* < 0.05. **F,** HCC4017 control or FCHSD2 siRNA-treated cells were stimulated with EGF (100 ng/ml) in the presence of cycloheximide (40 µg/ml) and incubated for the indicated times at 37°C. **G,** Quantification of EGFR/ERK intensity ratios in the cells as described in **F**. Percentage of degraded EGFR was calculated relative to the initial amount. Data represent mean ± SEM (*n* = 3). Two-tailed Student’s *t* tests were used to assess statistical significance. \**P* < 0.05.

FCHSD2 KD also reduced the efficiency of EGFR recycling (Fig. 1C; EV Fig. 2A and 2B). To further explore which stage in the process was disrupted, we followed p-EGFR trafficking using immunofluorescence and detected the accumulation of active EGFR in LAMP1-positive late endosome/lysosomes upon FCHSD2 KD (Fig. 1D and E; EV Fig. 2C and 2D). In accordance with these results, FCHSD2 depletion enhanced the rate of EGFR degradation following EGF stimulation (Fig. 1F and G). These findings reveal additional roles for FCHSD2 in endocytic recycling and in controlling EGFR trafficking and degradation.

**Figure 2.**
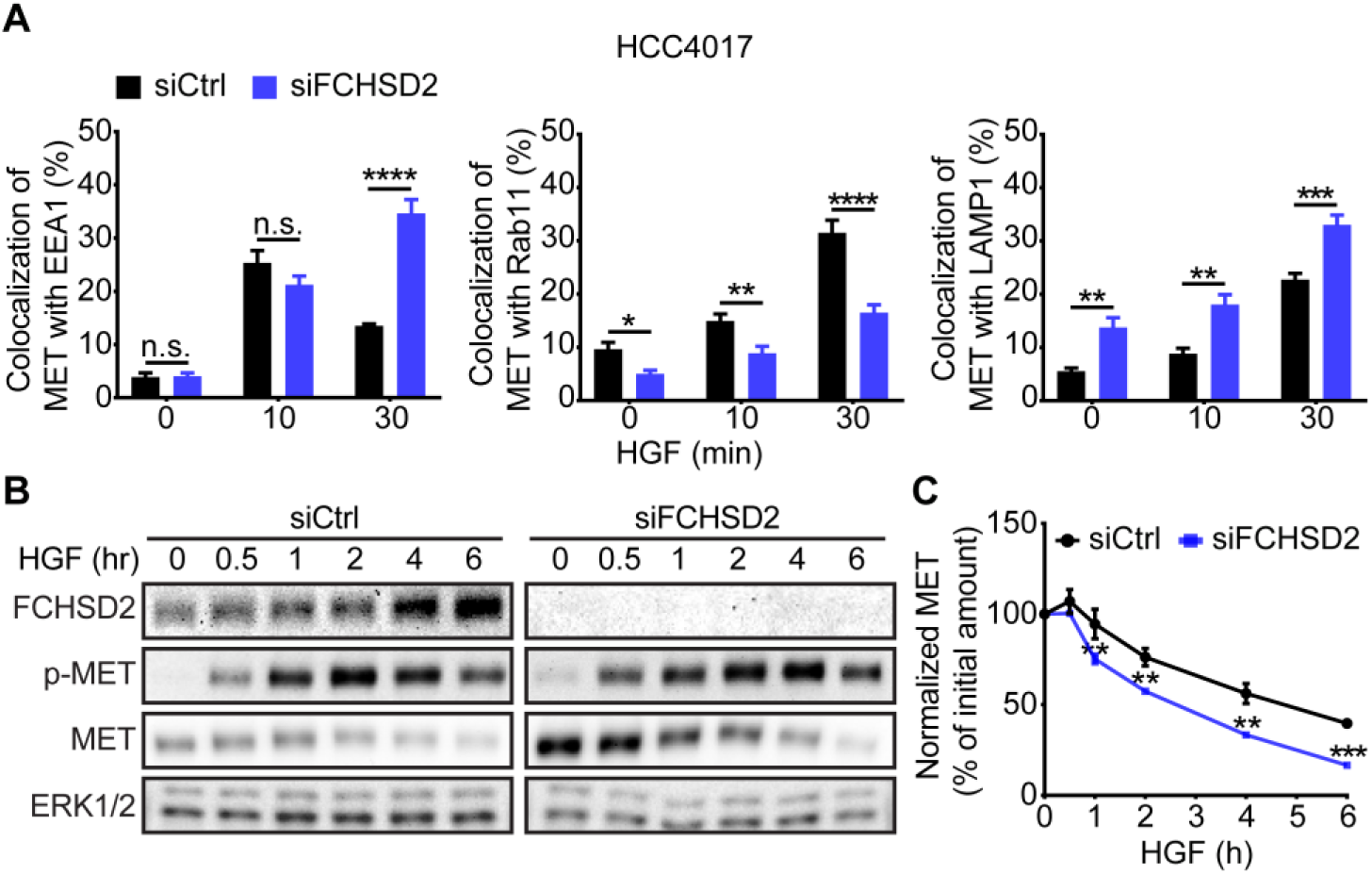
FCHSD2 depletion alters the trafficking of MET receptor. **A,** Quantification of the colocalization of MET with EEA1, Rab11 or LAMP1 immunofluorescence staining in control or FCHSD2 siRNA-treated HCC4017 cells. Cells were incubated with 1 μg/ml HGF for 30 min at 4°C, washed, and reincubated at 37°C for the indicated times. Data were obtained from at least 40 cells in total/condition and represent mean ± SEM. Two-tailed Student’s *t* tests were used to assess statistical significance. n.s., not significant, \**P* < 0.05, \*\**P* < 0.005, \*\*\**P* < 0.0005, \*\*\*\**P* < 0.00005. Representative confocal images are shown in EV Fig. 3 and 4. **B,** HCC4017 control or FCHSD2 siRNA-treated cells were stimulated with HGF (100 ng/ml) in the presence of cycloheximide (40 µg/ml) and incubated at 37°C for the indicated times. **C,** Quantification of MET/ERK intensity ratios in the cells as described in **B**. Percentage of degraded MET was calculated relative to the initial amount. Data represent mean ± SEM (*n* = 3). Two-tailed Student’s *t* tests were used to assess statistical significance. \*\**P* < 0.005, \*\*\**P* < 0.0005.

### FCHSD2 directs the endocytic trafficking of MET receptor in NSCLC cells

We have shown that FCHSD2 regulates endocytosis of TfnR and EGFR and that it negatively regulates EGFR signaling from the cell surface (Xiao et al., 2018). Less studied, but also highly associated with NSCLC progression is the RTK, MET and its ligand HGF (hepatocyte growth factor). Moreover, it has been established that MET signaling requires endocytosis (Barrow-McGee & Kermorgant, 2014, Joffre, Barrow et al., 2011) and that it differentially signals from early vs. late endosomes (Menard, Parker et al., 2014). Given the role of FCHSD2 in endosomal recycling, we speculate that FCHSD2 also functions in regulating the endocytic trafficking of MET. To test this hypothesis, we performed immunofluorescence to measure MET trafficking after HGF stimulation, using antibodies against MET, as well as different endosomal proteins (i.e. EEA1, Rab11 and LAMP1 that mark, respectively, early endosomes, recycling endosomes and late endosomes/lysosomes) (EV Fig. 3 and Fig. 4).

**Figure 3.**
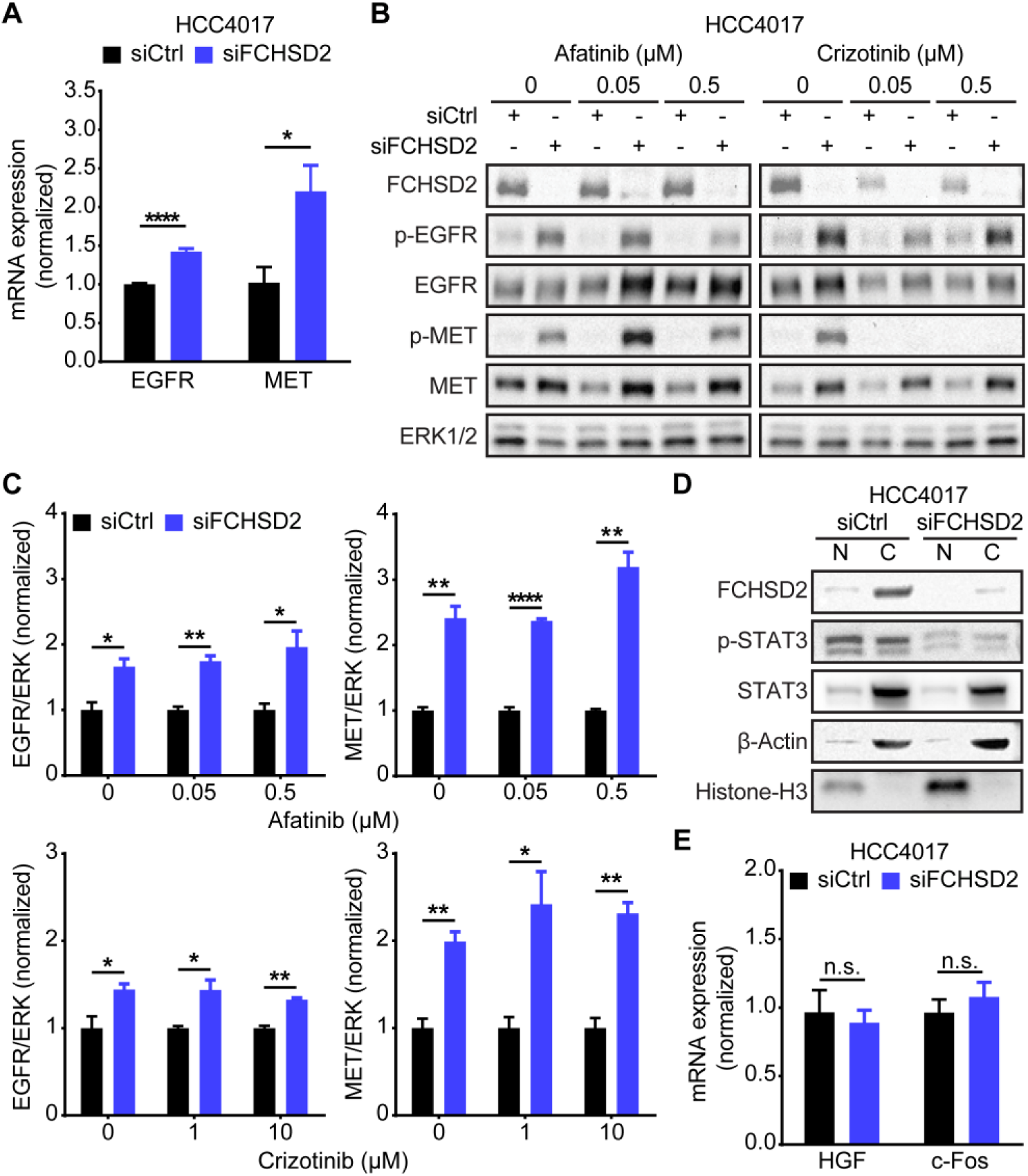
FCHSD2 depletion-induced upregulation of the RTKs is independent of their activities. **A,** FCHSD2 knockdown increases the transcription of *EGFR* and *MET* mRNA. All data were normalized to siCtrl and represent mean ± SEM (*n* = 3). Two-tailed Student’s *t* tests were used to assess statistical significance. \**P* < 0.05, \*\*\*\**P* < 0.00005. **B,** HCC4017 control or FCHSD2 siRNA-treated cells were incubated with EGFR inhibitor (Afatinib) or MET inhibitor (Crizotinib) at the indicated concentration for 24 h. **C,** Quantification of EGFR/ERK or MET/ERK intensity ratios in the cells as described in **B**. All data were normalized to siCtrl and represent mean ± SEM (*n* = 3). Two-tailed Student’s *t* tests were used to assess statistical significance. \**P* < 0.05, \*\**P* < 0.005, \*\*\*\**P* < 0.00005. **D,** Knockdown of FCHSD2 did not enhance translocation of phospho-STAT3 into the nucleus. Cell lysates from control or FCHSD2 siRNA-treated HCC4017 cells were subjected to fractionation. N, nuclear fraction. C, cytoplasmic fraction. **E,** Loss of FCHSD2 did not increase the transcription of phospho-STAT3 target genes, *HGF* and *c-Fos*. All data were normalized to siCtrl and represent mean ± SEM (*n* = 3). Two-tailed Student’s *t* tests were used to assess statistical significance. n.s., not significant.

**Figure 4.**
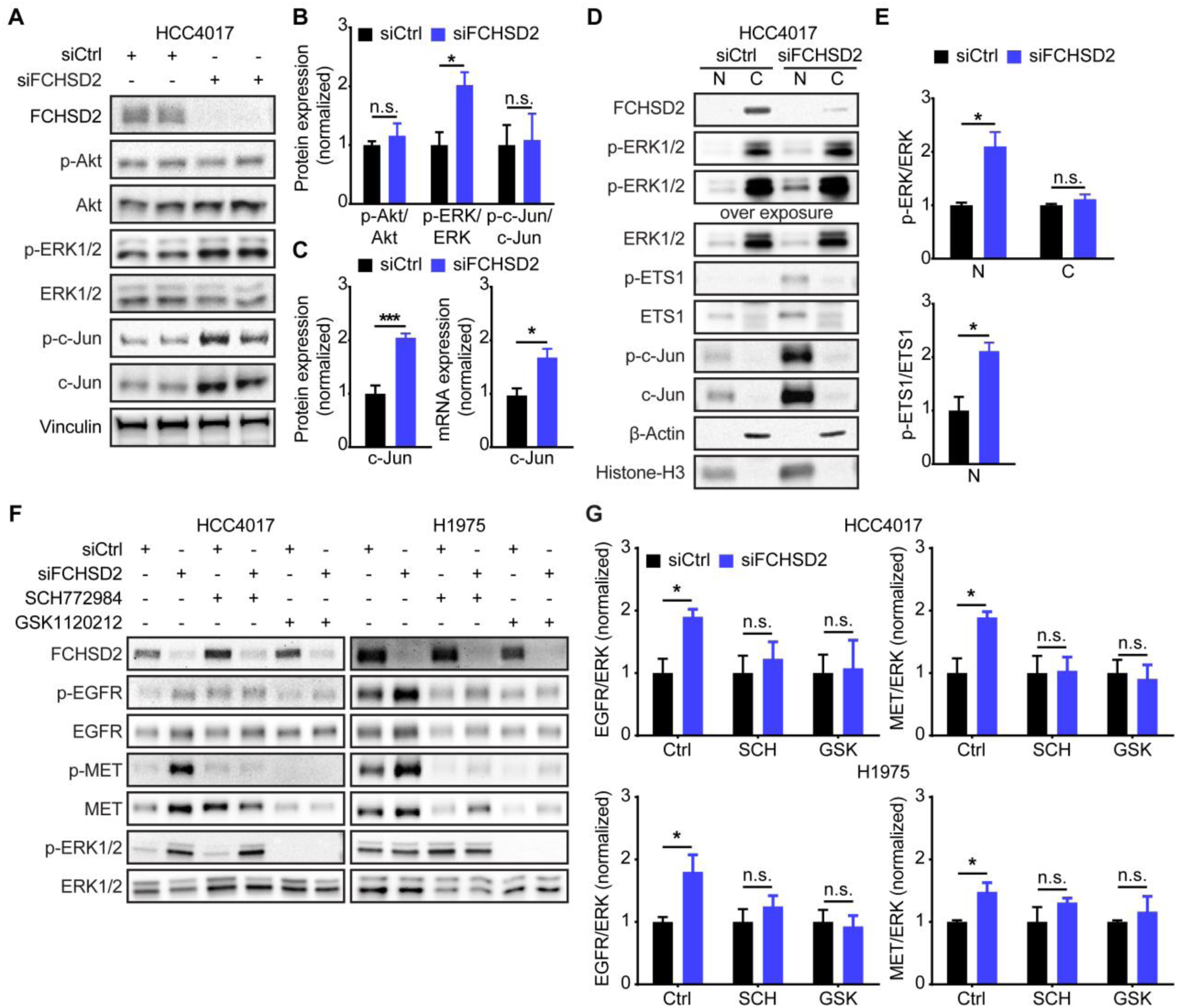
ERK1/2 activity is essential for the RTK upregulation induced by FCHSD2 depletion. **A,** Loss of FCHSD2 enhances ERK1/2 activity and c-Jun expression, but not Akt activity in HCC4017 cells. **B,** Quantification of signaling activities in the cells as described in **A**. All data were normalized to siCtrl and represent mean ± SEM (*n* = 3). Two-tailed Student’s *t* tests were used to assess statistical significance. n.s., not significant, \**P* < 0.05. **C,** Knockdown of FCHSD2 increases c-Jun protein expression and mRNA transcription. The protein expression was determined by quantification of c-Jun/Vinculin intensity ratios in the cells as described in **A**. All data were normalized to siCtrl and represent mean ± SEM (*n* = 3). Two-tailed Student’s *t* tests were used to assess statistical significance. \**P* < 0.05, \*\*\**P* < 0.0005. **D,** Knockdown of FCHSD2 specifically enhances nuclear p-ERK1/2, p-ETS1 and c-Jun levels in HCC4017 cells. Cell lysates from control or FCHSD2 siRNA-treated HCC4017 cells were subjected to fractionation. N, nuclear fraction. C, cytoplasmic fraction. **E,** Quantification of p-ERK/ERK and p-ETS1/ETS1 intensity ratios in the cells as described in **D**. All data were normalized to siCtrl and represent mean ± SEM (*n* = 3). Two-tailed Student’s *t* tests were used to assess statistical significance. n.s., not significant, \**P* < 0.05. **F,** ERK or MEK inhibition disrupts the RTK upregulation induced by FCHSD2 knockdown. ERK1/2 inhibitor (SCH772984, 1 μM) or MEK1/2 inhibitor (GSK1120212, 1 μM) was used to treat control or FCHSD2 siRNA-treated HCC4017 or H1975 cells for 72 h. **G,** Quantification of EGFR/ERK or MET/ERK intensity ratios in the cells as described in **F**. All data were normalized to siCtrl and represent mean ± SEM (*n* = 3). Two-tailed Student’s *t* tests were used to assess statistical significance. n.s., not significant, \**P* < 0.05.

Loss of FCHSD2 resulted in the accumulation of MET in early and late endosomes, with a corresponding decrease in colocalization with recycling endosomes (Fig. 2A). Additionally, FCHSD2 KD increased MET degradation following HGF stimulation (Fig. 2B and C). These data demonstrate that FCHSD2 also regulates trafficking and degradation of MET.

### FCHSD2 KD-induced upregulation of RTKs is independent of their activities

Paradoxically, we also noted that despite decreased recycling and enhanced degradation of the RTKs following FCHSD2 KD, the steady-state levels of EGFR and MET were higher in the FCHSD2 deficient cells (Fig. 1F and Fig. 2B). Unexpectedly, FCHSD2 KD resulted in increased levels of both *EGFR* and *MET* mRNA (Fig. 3A), consistent with the increased expression seen at the protein level.

According to previous studies, the transfer of active RTKs to perinuclear endosomes triggers the juxtanuclear activation of a weak STAT3 signal that leads to the required threshold of phosphorylation for nuclear translocation (Kermorgant & Parker, 2008, Miaczynska, 2013). In turn, accumulation of nuclear p-STAT3 promotes the transcription of *HGF* and *c-Fos* (Carpenter & Lo, 2014), leading to upregulation of EGFR (Johnson, Murphy et al., 2000) and MET (Anastasi, Giordano et al., 1997, Boccaccio & Comoglio, 2006). To test whether this signaling pathway accounted for our findings, we treated cells with an EGFR inhibitor (afatinib) or a MET inhibitor (crizotinib). However, neither inhibition affected the upregulation of RTKs in the FCHSD2 depleted cells (Fig. 3B and C; EV Fig. 5). Moreover, FCHSD2 KD decreased the level of p-STAT3 (Fig. 3D) and was unable to trigger the transcription of STAT3 target genes, *HGF* and *c-Fos* (Fig. 3E).

**Figure 5.**
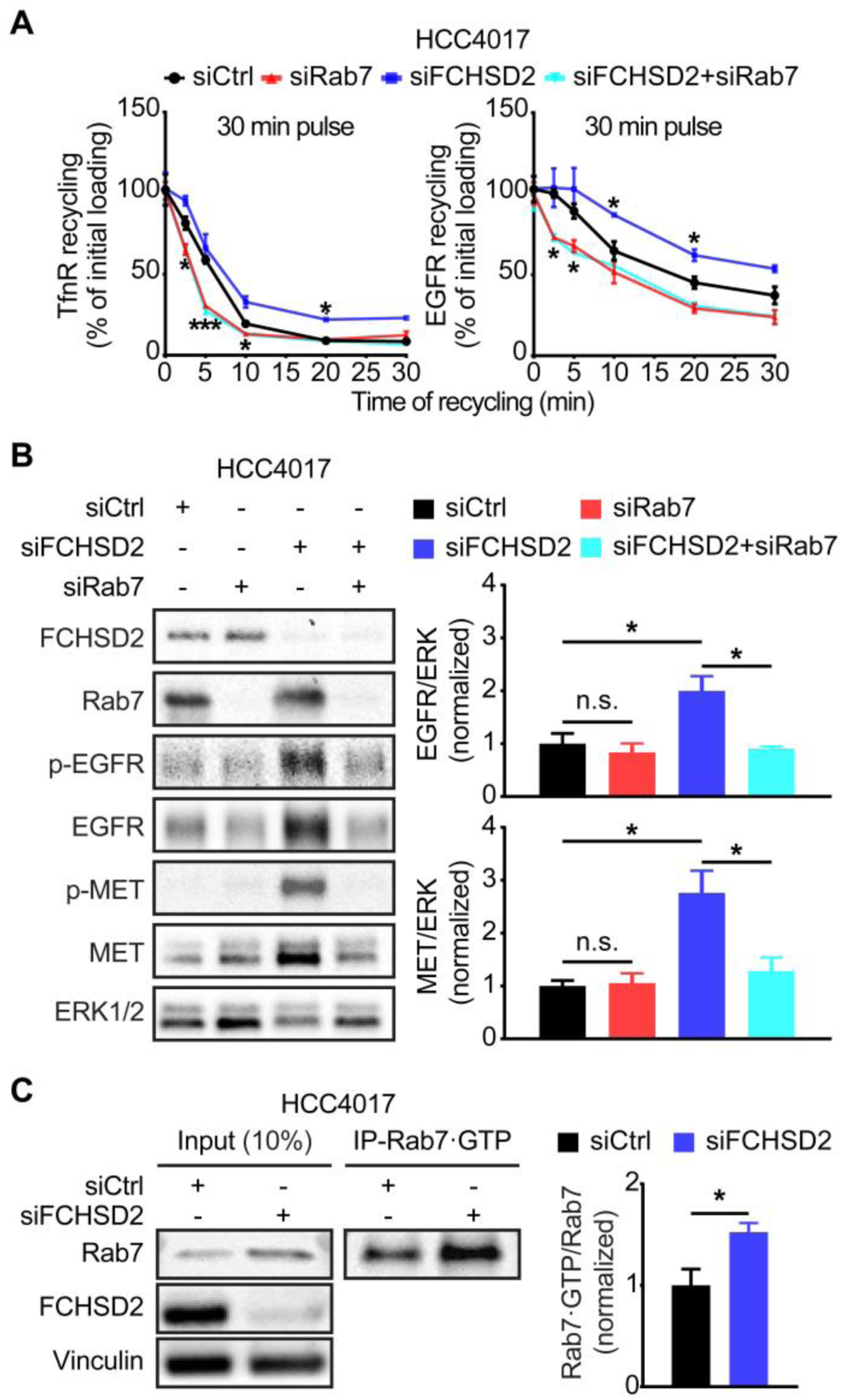
Rab7 is required for FCHSD2 depletion-induced upregulation of the RTKs. **A,** Endocytic recycling of TfnR or EGFR was measured in control, Rab7, FCHSD2 or both siRNA-treated HCC4017 cells. Cells were pulsed for 30 min with 10 μg/ml biotinylated Tfn or 20 ng/ml biotinylated EGF, stripped, and reincubated at 37°C for the indicated times before measuring the remaining intracellular Tfn or EGF. Percentage of recycled biotinylated Tfn or EGF was calculated relative to the initial loading. Data represent mean ± SEM (*n* = 3). Two-tailed Student’s *t* tests were used to assess statistical significance versus siCtrl. \**P* < 0.05, \*\*\**P* < 0.0005. **B,** Rab7 knockdown abolishes the RTK upregulation induced by FCHSD2 depletion. Quantification of EGFR/ERK or MET/ERK intensity ratios in the cells was measured. All data were normalized to siCtrl and represent mean ± SEM (*n* = 3). Two-tailed Student’s *t* tests were used to assess statistical significance. n.s., not significant, \**P* < 0.05. **C,** FCHSD2 depletion promotes the activity of Rab7. Cell lysates from control or FCHSD2 siRNA-treated HCC4017 cells were immunoprecipitated with anti-Rab7·GTP (active form of Rab7) antibody. The indicated proteins were detected. Quantification of Rab7·GTP/input Rab7 intensity ratios in the cells was measured. All data were normalized to siCtrl and represent mean ± SEM (*n* = 3). Two-tailed Student’s *t* tests were used to assess statistical significance. \**P* < 0.05.

### ERK1/2 activity is responsible for the FCHSD2 KD-induced RTK upregulation

Having ruled out STAT3 signaling and indeed, activities of the RTKs themselves, we next looked for alterations in steady-state activity of other signaling pathways that might account for the upregulation of ERK and MET upon FCHSD2 KD. We observed that FCHSD2 depletion specifically increased ERK1/2 activity, but not Akt activity even at steady-state in HCC4017 cells (Fig. 4A and B). Notably, FCHSD2 KD significantly increased the expression of c-Jun and p-c-Jun, although the ratio of p-c-Jun/c-Jun was unaffected (Fig. 4A-C). Constitutive activation of ERK1/2 signaling induces *c-Jun* transcription and sustains c-Jun stability and activity (Lopez-Bergami, Huang et al., 2007); correspondingly, FCHSD2 KD enhanced the transcription of *c-Jun* mRNA (Fig. 4C). Further, loss of FCHSD2 specifically increased ERK1/2 activity in the nucleus, while the ratio of pERK/ERK in the cytoplasm remained unchanged (Fig. 4D and E). In addition to enhancing c-Jun expression, the accumulation of nuclear p-ERK1/2 in FCHSD2-depleted cells promoted activity of the ERK1/2 target, ETS1 (Plotnikov, Zehorai et al., 2011) (Fig. 4D and E). Both c-Jun and ETS1 are known transcription factors for *EGFR* (Johnson et al., 2000) and *MET* (Boccaccio & Comoglio, 2006, Gambarotta, Boccaccio et al., 1996), accounting for the observed increase in transcription of *EGFR* and *MET* mRNA after FCHSD2 KD (Fig. 3A).

To directly test whether ERK1/2 activity is required for increased expression of the RTKs in FCHSD2 KD cells, we used an ERK1/2 kinase inhibitor (SCH772984) and an inhibitor targeting the essential upstream kinase, MEK1/2 (GSK1120212). As predicted, both ERK1/2 and MEK1/2 inhibition disrupted the upregulation of EGFR and MET receptor in HCC4017 and H1975 cells (Fig. 4F and G). Together, these results suggest that increased ERK1/2 activity in the nucleus is essential for the effects of FCHSD2 depletion in NSCLC cells.

### Rab7 is essential for the effects of FCHSD2 KD on RTK expression

Previous studies have shown that translocation of active ERK1/2 to the nucleus requires the recruitment of MEK1 to Rab7-positive endosomes, where MEK1 activates ERK1/2 signaling from late/perinuclear endosome compartments (Nada, Hondo et al., 2009). In addition, Rab7 supports endosome maturation and promotes endocytic trafficking toward late endosomes rather than to recycling endosome compartments (Langemeyer, Frohlich et al., 2018). In agreement with previous research, Rab7 KD increased the rate of recycling of TfnR and EGFR (Fig. 5A). Strikingly, there was no difference between the effects of Rab7 KD alone and the depletion of both FCHSD2 and Rab7 on the TfnR and EGFR recycling (Fig. 5A), indicating that the FCHSD2 KD-induced phenotype depends on the function of Rab7. Given that the activities of MEK1/2 and ERK1/2 are necessary for the effects of FCHSD2 KD, we further tested the consequences of Rab7 depletion in the cells. Importantly, Rab7 KD abolished the upregulation of EGFR and MET induced by FCHSD2 depletion (Fig. 5B).

Rab7 is a small GTPase, whose function is determined by its expression and activity, regulated by a switch between active GTP-bound (Rab7·GTP) and inactive GDP-bound (Rab7·GDP) states (Langemeyer et al., 2018). To assess the effect of FCHSD2 KD on Rab7 activity, we immunoprecipitated active Rab7 using an antibody specifically recognizing the Rab7·GTP in cancer cells. We found that there were higher levels of active Rab7 in the FCHSD2-deficient cells (Fig. 5C). These data suggest that FCHSD2 controls expression and trafficking of the RTKs by negatively regulating Rab7. Thus, FCHSD2 and Rab7 play antagonistic roles in regulating endosomal trafficking.

### FCHSD2 and Rab7 differentially effect lung cancer progression

Our studies have revealed a multifaceted function for FCHSD2 in the crosstalk between endocytic trafficking and oncogenic signaling in NSCLC cells (Fig. 6A). We previously showed that FCHSD2 has a cancer cell-specific function in positively regulating CME downstream of ERK1/2 activity. Here we show that FCHSD2 plays a general role in regulating endosomal trafficking through adversely modulating Rab7 activity. Together these activities establish FCHSD2 as a key regulator in oncogenic ERK1/2 signaling outcome by controlling the trafficking and expression of EGFR and MET. Given that FCHSD2 KD dramatically increased the proliferation and the migration activities of NSCLC cells (Xiao et al., 2018), FCHSD2 may function as a negative regulator for human lung tumor growth. In contrast, Rab7 is thought to favor lung cancer progression.

**Figure 6.**
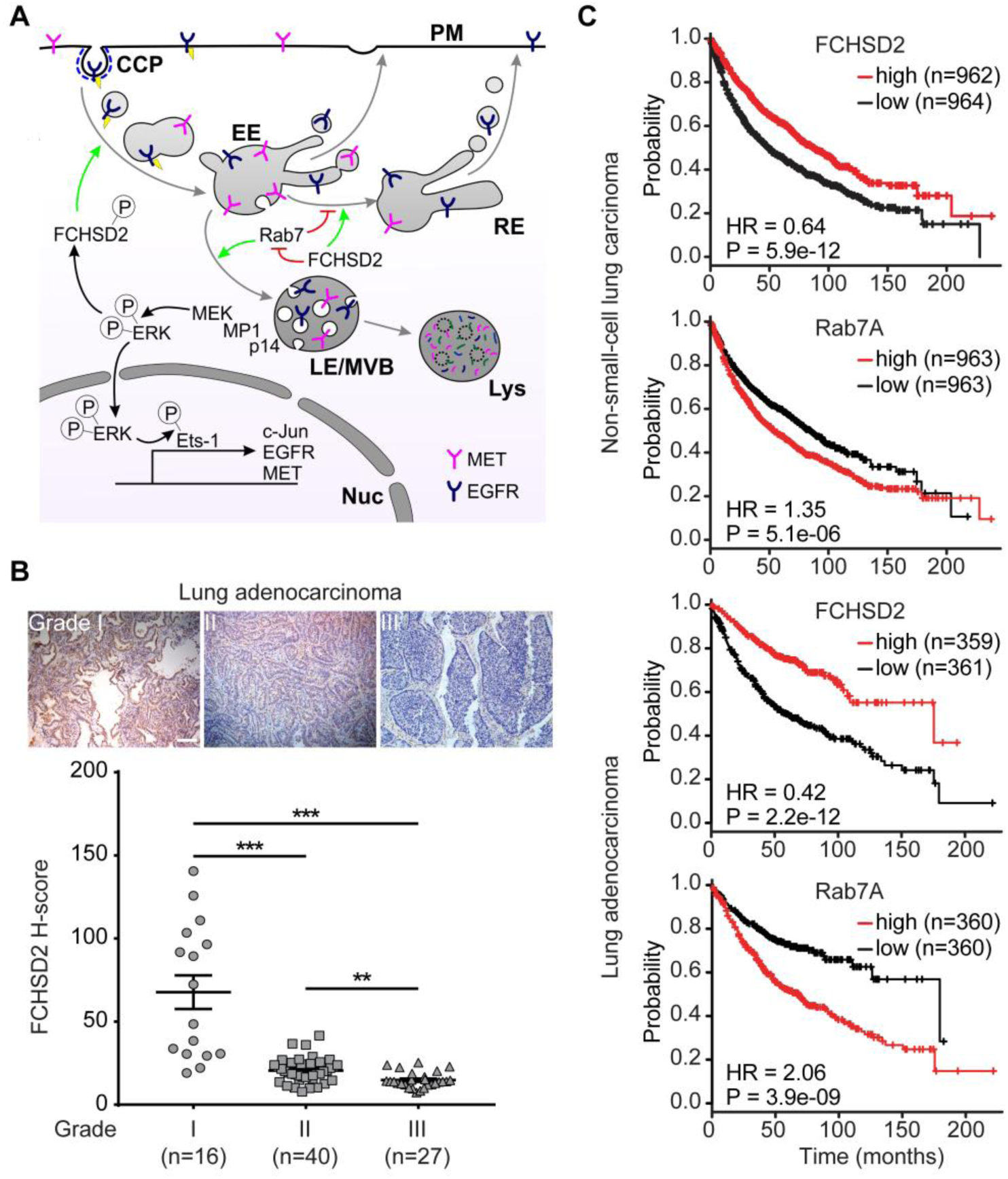
FCHSD2 and Rab7 reciprocally regulate endocytic trafficking and lung cancer progression. **A,** FCHSD2 regulates multiple steps in endocytic trafficking. We previously showed that activation of FCHSD2 downstream of ERK1/2 increases the rate of clathrin-coated pits (CCP) initiation and CME in NSCLC cells. Here we report that FCHSD2 also increases the rate of RTK trafficking from early endosomes (EE) to recycling endosomes (RE) and negatively regulates Rab7 activity, maturation of late endosomes/multivesicular bodies (LE/MVB) and trafficking to lysosomes (Lys). Together these activities of FCHSD2 increase the flux of RTKs through early endocytic pathways and thus altering their downstream signaling. Loss of FCHSD2 results in the accumulation of RTKs in late endosomes/lysosomes, increases levels of activated ERK1/2 in the nucleus and enhances transcription and expression of c-Jun, EGFR and MET. **B,** Immunohistochemistry images and quantification (expressed a H-score) of FCSHD2 staining in representative lung tumor tissues. Scale bar, 100 μm. **C,** Kaplan-Meier survival analysis of NSCLC or lung adenocarcinoma patients was performed in FCHSD2 or Rab7 high- and low-expression cohorts.

To investigate the correlation between FCHSD2 activity and lung tumor progression, we directly measured the protein expression level of FCHSD2 in tumor tissues from lung adenocarcinoma patients. As expected from our previous *in vitro* findings, FCHSD2 expression is gradually decreased in higher grades of lung adenocarcinoma tumors (Fig. 6B).

Finally, we examined the relationship between FCHSD2 and Rab7 and NSCLC patient survival by mining clinical data. Patients with relatively high FCHSD2 expression had significantly better survival rates than those in the low-expression group (Fig. 6C). In contrast, Rab7 expression had the opposite correlation with patient survival rates (Fig. 6C). Notably, the correlation between the expression of FCHSD2 or Rab7 with survival rates was more prominent in lung adenocarcinoma patients (Fig. 6C). These findings suggest that FCHSD2 functions as a negative regulator of Rab7 and controls lung cancer aggressiveness.

## Discussion

Endocytic trafficking regulates the expression and activity of RTKs and modulates their downstream signaling to maintain cell homeostasis (Antonescu, McGraw et al., 2014). We previously reported that activation of FCHSD2 by ERK1/2 phosphorylation increases the rate of TfnR and EGFR internalization by CME and suppresses signaling from cell surface EGFRs, specifically in cancer cells (Xiao et al., 2018). Here we show that FCHSD2, like its *Drosophila* orthologue *Nwk* (Rodal et al., 2011) also enhances recycling of internalized RTKs and reduces their trafficking to late endosomes/lysosomes. This activity is independent of ERK activation and involves the negative regulation of Rab7. Together these FCHSD2-dependent changes enhance the rate of trafficking of RTKs through the early and recycling endocytic pathways. As a result, their signaling pathways, in particular those downstream of ERK1/2 nuclear signaling, are suppressed resulting in decreased proliferation and reduced cell migration. Our mechanistic studies of FCSHD2 function in NSCLC cell lines were consistent with clinical databases showing that high levels of FCHSD2 expression correlate with improved survival rates, especially among lung adenocarcinoma patients, whose cancers are frequently driven by oncogenic mutations that activate MAP kinase signaling (Dogan, Shen et al., 2012, Ferrer, Zugazagoitia et al., 2018). Together, these data suggest that the regulation of early endocytic trafficking by FCHSD2 functions to suppress signaling downstream of activated RTKs potentially as a means to maintain cell homeostasis.

Overexpressed RTKs are a common feature among different types of cancers and widely considered favorable for tumor progression (Maegawa, Arao et al., 2009). In particular, MET, the HGF receptor, is upregulated in ∼50% of NSCLC, particularly in lung adenocarcinomas (72.3%) (Ichimura, Maeshima et al., 1996). Hyperactivity of MET and its dependent invasive growth signals is a general feature of highly aggressive tumors and associated with poor survival (Comoglio, Giordano et al., 2008). Moreover, the activation of MET and its downstream signaling is dependent on trafficking through both early and late endosomes (Joffre et al., 2011, Trusolino, Bertotti et al., 2010). FCHSD2 depletion resulted in both increased expression of MET and decreased recycling leading to the accumulation of MET in both early and late endosome/lysosomes compartments. These FCHSD2-dependent changes in MET expression and trafficking, are consistent with our immunohistochemistry studies showing that FCHSD2 expression levels inversely correlated with more advanced stages of lung adenocarcinomas.

FCHSD2 KD led to increased steady-state expression levels of the oncogenes MET, EGFR and c-Jun, as well as an increase in the steady-state activation of ERK1/2 specifically in the nucleus. Paradoxically, these changes were not dependent on either MET or EGFR kinase activities, but required ERK1/2 activity. A previous study showed that KRas can be constitutively internalized via CME and activated on Rab7-positive late endosomes (Lu, Tebar et al., 2009). There the late endosome-associated adaptor p14 and the scaffolding protein MP1 tether MEK and ERK1/2 for downstream activation of ERK1/2 (Wunderlich, Fialka et al., 2001). We speculate that, even if inactive, the high local concentrations of EGFR and MET that accumulate on late endosomes after FCHSD2 KD may be sufficient to recruit mSOS and activate KRas. Interestingly, basal Ras activation has also been reported in Neimann-Pick C fibroblasts where late endosomal trafficking is perturbed (Corey & Kelley, 2007).

Rab7 is a ubiquitously expressed member of the Rab family of small GTPases localized to late endosomal compartments and plays a vital role in endosomal membrane traffic (Guerra & Bucci, 2016). Specifically, Rab7 mediates the maturation of early endosomes into late endosomes, fusion of late endosomes with lysosomes in the perinuclear region and lysosomal biogenesis (Langemeyer et al., 2018). Here we show that the effects of FCHSD2 KD on endosomal trafficking and consequent upregulation of EGFR and MET expression are dependent on Rab7 and that FCHSD2 appears to negatively regulate Rab7 activation. These findings show that Rab7 and FCHSD2 functions in endosomal trafficking are antagonistic. Consistent with this, we report here that the expression of Rab7 is negatively correlated with NSCLC patient survival, which is opposite to the positive correlation between FCHSD2 expression and patient survival, especially in lung adenocarcinomas. That these two proteins converge on regulating trafficking between early endosomes, recycling endosomes and late endosomes suggests an important role for endosomal trafficking in regulating signaling in cancer cells and tumor progression.

Collectively, our findings define an endocytic trafficking pathway regulated by FCHSD2 and Rab7 that functions to control RTK expression, oncogenic signal transduction and NSCLC progression. Knowledge of these trafficking pathways and their (dys)regulation during cancer progression could help to identify potential new therapeutic targets for the prevention of aggressive cancers and/or prognostic indicators that can guide lung cancer treatment.

## Material and Methods

### Cell culture and chemicals

HCC4017 (KRas^G12C^, EGFR^WT^) and H1975 (KRas^WT^, EGFR^T790M/L858R^) NSCLC cells (from John Minna, UT Southwestern Medical Center, Dallas) were grown in RPMI 1640 (Thermo Fisher Scientific) supplemented with 10% (vol/vol) FCS (HyClone). ARPE-19 cells (from ATCC) were cultivated in DMEM/F12 (Thermo Fisher Scientific) supplemented with 10% (vol/vol) FCS. The recombinant human EGF used in this study was from Thermo Fisher Scientific and the recombinant human HGF was generously provided by Drs. Emiko Uchikawa and Xiaochen Bai (UT Southwestern Medical Center, Dallas). The cycloheximide was from MilliporeSigma. The EGFR inhibitor Afatinib, the MET inhibitor Crizotinib and the ERK inhibitor SCH772984 were from Selleck Chemicals. The MEK inhibitor GSK1120212 was from MedChemExpress.

### RNA interference

Cells were treated with the siRNA pool targeting FCHSD2 (#J-021240-10, J-021240-11, J-021240-12, Dharmacon) or Rab7 (LU-010388-00, Dharmacon) using RNAiMAX (Thermo Fisher Scientific) to silence the endogenous protein. Briefly, 50 nM of the indicated siRNA pool and 6.5 μl of RNAiMAX reagent were added in 1 ml of OptiMEM (Thermo Fisher Scientific) to each well of a 6-well plate and incubated for 20 min at room temperature. Cells were resuspended in 1 ml of culture medium, seeded in each well of a 6-well plate at 20-30% confluency containing the mixed siRNA-lipid complex and incubated for 48-72 h, followed by experiments. The AllStars Negative siRNA non-targeting sequence was purchased from Qiagen (#SI03650318).

### Western blotting and analyses

Cells cultured in each well of a 6-well plate at 80% confluency were washed three times with PBS and harvested/resuspended in 150–200 μl of 2× Laemmli buffer (Bio-Rad). The cell lysate was boiled for 10 min and loaded onto an SDS gel. After transferring to a nitrocellulose membrane (Bio-Rad), membranes were blocked with 5% milk in TBST buffer and were probed with antibodies diluted in 5% BSA in TBST buffer against the following proteins: FCHSD2 (#PA5-58432, 1:500, Thermo Fisher Scientific), Rab11 (#5589S, 1:1000, Cell Signaling), Rab7 (#9367S, 1:1000, Cell Signaling), p-EGFR Y1068 (#3777S, 1:1000, Cell Signaling), EGFR (#4267S, 1:1000, Cell Signaling), p-MET Y1234/1235 (#3077S, 1:1000, Cell Signaling), MET (#8198S, 1:1000, Cell Signaling), p-Akt S473 (#4060L, 1:1000, Cell Signaling), Akt (#9272S, 1:1000, Cell Signaling), p-ERK1/2 T202/Y204 (#4370S, 1:1000, Cell Signaling), ERK1/2 (#4695S, 1:1000, Cell Signaling), p-STAT3 Y705 (#4370S, 1:1000, Cell Signaling), STAT3 (#9139S, 1:1000, Cell Signaling), p-c-Jun S63 (#9261S, 1:1000, Cell Signaling), c-Jun (#9165S, 1:1000, Cell Signaling), p-ETS1 T38 (#ab59179, 1:1000, Abcam), ETS1 (#14069S, 1:1000, Cell Signaling), β-Actin (#sc-47778, 1:2500, Santa Cruz), Histone-H3 (#4499S, 1:2000, Cell Signaling) and Vinculin (#V9131, 1:1000, MilliporeSigma). Horseradish peroxidase (HRP)-conjugated secondary antibodies (#G21234 and # G21040, 1:2000, Thermo Fisher Scientific) were used according to the manufacturers’ instructions. Quantitative analysis was performed by using ImageJ software (NIH).

For EGF- or HGF-induced degradation of EGFR or MET receptors, after siRNA transfection, the cells (5×10^5^) were seeded in each well of a 6-well plate containing RPMI 1640 with 10% FCS. Eight hours after seeding, cells were washed three times with PBS and starved in RPMI 1640 without FCS for 16 h. The cells then were untreated or treated with 100 ng/ml of EGF or HGF in the presence of cycloheximide (40 µg/ml) for the indicated times. After the stimulation, cells were washed three times with PBS and harvested/resuspended in 150–200 μl of 2× Laemmli buffer, and the cell lysates were subjected to Western blotting and image analysis as described above.

### Endocytic recycling assay

TfnR recycling assays were performed using biotinylated Tfn, which was conjugated at a 7:1 molar ratio with the cleavable EZ-Link Sulfo-NHS-SS-Biotin (#A39258, Thermo Fisher Scientific) according to manufacturer’s instructions. For EGFR recycling, non-cleavable biotinylated EGF (#E3477, Thermo Fisher Scientific) was used for assays. TfnR and EGFR recycling assays were performed as previously described (Chen et al., 2017). In brief, cells were grown overnight in gelatin-coated 96-well plates at a density of 3×10^4^ cells/well and incubated with 10 μg/ml biotinylated Tfn or 20 ng/ml biotinylated EGF in assay buffer (PBS^4+^: PBS supplemented with 1 mM MgCl_2_, 1 mM CaCl_2_, 5 mM glucose and 0.2% bovine serum albumin) for the 10 or 30 min pulse at 37°C. Cells were then immediately cooled down (4°C) to stop internalization. The remaining surface-bound biotinylated Tfn was cleaved by incubation with 10 mM Tris (2-carboxyethyl) phosphine (TCEP) in assay buffer for 30 min at 4°C. The surface-bound biotinylated EGF was removed by acid wash (0.2 M acetic acid, 0.2 M NaCl, pH 2.5) at 4°C. For TfnR recycling assays using ERK1/2 inhibitors (SCH772984), cells were incubated in the absence or presence of 10 µM of SCH772984 in the assay buffer containing 10 mM TCEP for 30 min at 4°C before recycling assays were performed in the continued absence or presence of the inhibitor. Cells were washed with cold PBS^4+^ buffer and then incubated in PBS^4+^ containing 2 mg/ml of holo-Tfn or 100 ng/ml of EGF and 10 mM of TCEP at 37°C for the indicated times. The recycled biotinylated Tfn or biotinylated EGF was removed from the cells by the acid wash step. Cells were then washed with cold PBS and then fixed in 4% paraformaldehyde (PFA) (Electron Microscopy Sciences) in PBS for 30 min and further permeabilized with 0.1% Triton X-100/PBS for 10 min. Remaining intracellular biotinylated ligands were assessed by streptavidin-POD (#11089153001, 1:10000, Roche) in Q-PBS, which contains 0.2% BSA (Equitech-Bio), 0.001% saponin (MilliporeSigma), and 0.01% glycine (MilliporeSigma). The reaction was further developed with OPD (MilliporeSigma), and then stopped by addition of 50 μl of 5 M of H_2_SO_4_. The absorbance was read at 490 nm (Biotek Synergy H1 Hybrid Reader). The decrease in intracellular biotinylated ligands (recycling) were represented as the percentage of the total internal pool of ligand internalized. Well-to-well variability in cell number was accounted for by normalizing the reading at 490 nm with a BCA readout at 562 nm.

### Immunofluorescence and confocal microscopy analyses

After siRNA transfection, the cells were washed three times with PBS and then starved in RPMI 1640 medium without FCS for 30 min at 37°C. Cells were incubated with 20 ng/ml EGF or 1 μg/ml HGF in RPMI 1640 medium for 30 min at 4°C, washed with cold PBS and then incubated in pre-warmed RPMI 1640 medium at 37°C for the indicated times. Cells were then washed with ice-cold PBS to stop chase, fixed with 4% (w/v) PFA for 30 min at 37°C and permeabilized using 0.05% saponin (w/v) (MilliporeSigma) for 10 min. Cells were blocked with Q-PBS and probed with antibodies against the following proteins: p-EGFR Y1068 (#3777S, 1:200, Cell Signaling), MET (#AF276, 1:100, R&D Systems), EEA1 (#610457, 1:100, BD Biosciences), Rab11 (#5589S, 1:50, Cell Signaling), and LAMP1 (#ab25245, 1:75, Abcam), according to the manufacturers’ instructions. AlexaFluor-conjugated secondary antibodies (#A-11036, A-11055, A-21206, A-21434, A-31571, Thermo Fisher Scientific) were used according to the manufacturers’ instructions. Fixed cells were mounted in PBS and imaged using a 60×, 1.49 NA APO objective (Nikon) mounted on a Nikon Ti-Eclipse inverted microscope coupled to an Andor Diskovery Spinning disk confocal/Borealis widefield illuminator equipped with an additional 1.8× tube lens (yielding a final magnification of 108×). The pinhole size was 50 µm. The percentages of colocalizations were determined using ImageJ software (NIH).

### Nuclear/cytosol fractionation

After siRNA transfection, the cells were subjected to fractionation using the Nuclear/Cytosol Fractionation Kit (K266-25, BioVision) according to the manufacturer’s instructions.

### Rab7 activation assay

After siRNA transfection, the cells were collected in ice-cold PBS containing protease and phosphatase inhibitor cocktails (Roche). The cells were used to measure Rab7 activation using the Rab7 Activation Assay Kit (#NEBB40025, NewEast Biosciences) according to the manufacturer’s instructions.

### Analysis of Kaplan-Meier Survival Data

NSCLC patient survival data was downloaded from the Kaplan Meier plotter database (Gyorffy, Surowiak et al., 2013). Analysis of NSCLC patients was performed in FCHSD2 or Rab7 high and low expression cohort. *P* value was calculated by logrank test (Gyorffy et al., 2013).

### Immunohistochemistry and image analyses

The human lung adenocarcinoma tissues were from US Biomax Inc (#LC641) and the UT Southwestern Tissue Resource, a shared resource at the Simmons Comprehensive Cancer Center. The tumors were classified according to the American Joint Committee on Cancer (AJCC) TNM system. The immunohistochemical staining of FCHSD2 (#PA5-58432, Thermo Fisher Scientific) was optimized and performed by the core facility. The immunohistochemical images were analyzed using the IHC Profiler, ImageJ software (NIH) as previously described (Varghese, Bukhari et al., 2014) to classify the intensities of staining. The immunoreactivity was determined by H-score, generated by adding the percentage of strong staining (3×), the percentage of moderate staining (2×) and the percentage of weak staining (1×) samples (Goulding, Pinder et al., 1995).

## Authors’ Contributions

**Conception and design:** G.Y. Xiao, S. L. Schmid

**Development of methodology:** G.Y. Xiao

**Acquisition of data (provided animals, acquired and managed patients, provided facilities, etc.):** G.Y. Xiao

**Analysis and interpretation of data (e.g., statistical analysis, biostatistics, computational analysis):** G.Y. Xiao

**Writing, review, and/or revision of the manuscript:** G.Y. Xiao, S. L. Schmid

**Administrative, technical, or material support (i.e., reporting or organizing data, constructing databases):** G.Y. Xiao, S. L. Schmid

**Study supervision:** S. L. Schmid

## Acknowledgments

We thank members of the Schmid lab for critically reading the manuscript, and especially Marcel Mettlen for his help preparing illustrations. We thank colleagues from Department of Biophysics: Drs. Emiko Uchikawa and Xiaochen Bai for kindly providing recombinant human HGF. We acknowledge the UT Southwestern Tissue Resource, a shared resource at the Simmons Comprehensive Cancer Center (supported in part by the National Cancer Institute under award number 5P30CA142543) for assistance with immunohistochemistry data analysis and interpretation. The work was supported by NIH grants R01 GM45455 and GM73165 to SLS.

---

**EV Figure 1.**
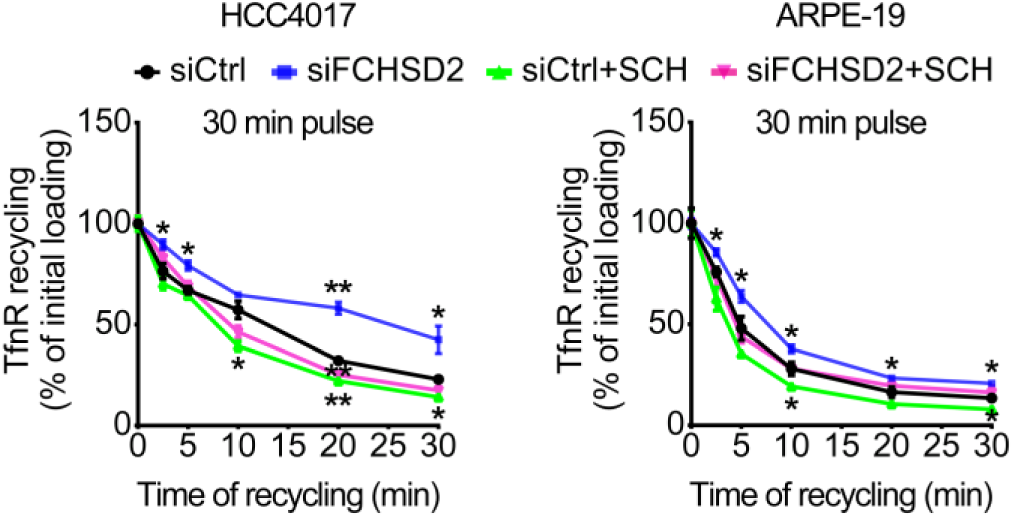
The effects of FCHSD2 depletion and ERK1/2 inhibition in TfnR endocytic recycling. Endocytic recycling of TfnR was measured in control or FCHSD2 siRNA-treated HCC4017 and ARPE-19 cells in the absence or presence of the ERK1/2 inhibitor SCH772984 (10 μM). Cells were pulsed for 30 min with 10 μg/ml biotinylated Tfn, stripped, and reincubated at 37°C for the indicated times before measuring the remaining intracellular Tfn. Percentage of recycled biotinylated Tfn was calculated relative to the initial loading. Data represent mean ± SEM (*n* = 3). Two-tailed Student’s *t* tests were used to assess statistical significance versus siCtrl. \**P* < 0.05, \*\**P* < 0.005.

**EV Figure 2.**
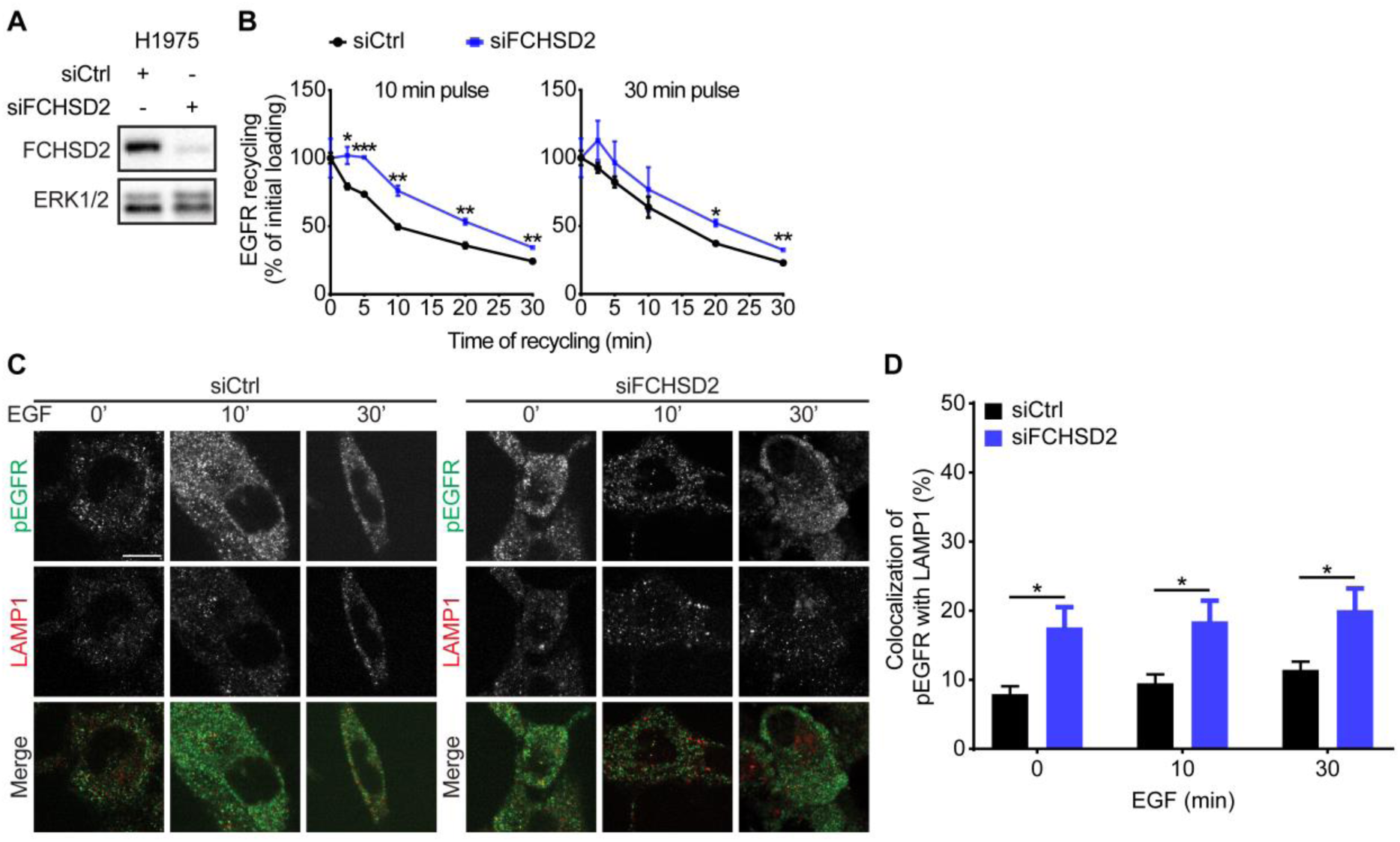
FCHSD2 regulates EGFR endocytic trafficking in H1975 cells. **A,** The knockdown of FCHSD2 in control or FCHSD2 siRNA-treated H1975 cells. **B,** Endocytic recycling of EGFR was measured in control or FCHSD2 siRNA-treated H1975 cells. Cells were pulsed for 10 min or 30 min with 20 ng/ml biotinylated EGF, stripped, and reincubated at 37°C for the indicated times before measuring the remaining intracellular EGF. Percentage of recycled EGF was calculated relative to the initial loading. Data represent mean ± SEM (*n* = 3). Two-tailed Student’s *t* tests were used to assess statistical significance. \**P* < 0.05, \*\**P* < 0.005, \*\*\**P* < 0.0005. **C,** Representative confocal images of pEGFR and LAMP1 immunofluorescence staining in control or FCHSD2 siRNA-treated H1975 cells. Cells were incubated with 20 ng/ml EGF for 30 min at 4°C, washed, and reincubated at 37°C for the indicated times. Scale bar, 12.5 μm. **D,** Colocalization of pEGFR and LAMP1 immunofluorescence staining in the cells as described in **C**. Data were obtained from at least 40 cells in total/condition and represent mean ± SEM. Two-tailed Student’s *t* tests were used to assess statistical significance. \**P* < 0.05.

**EV Figure 3.**
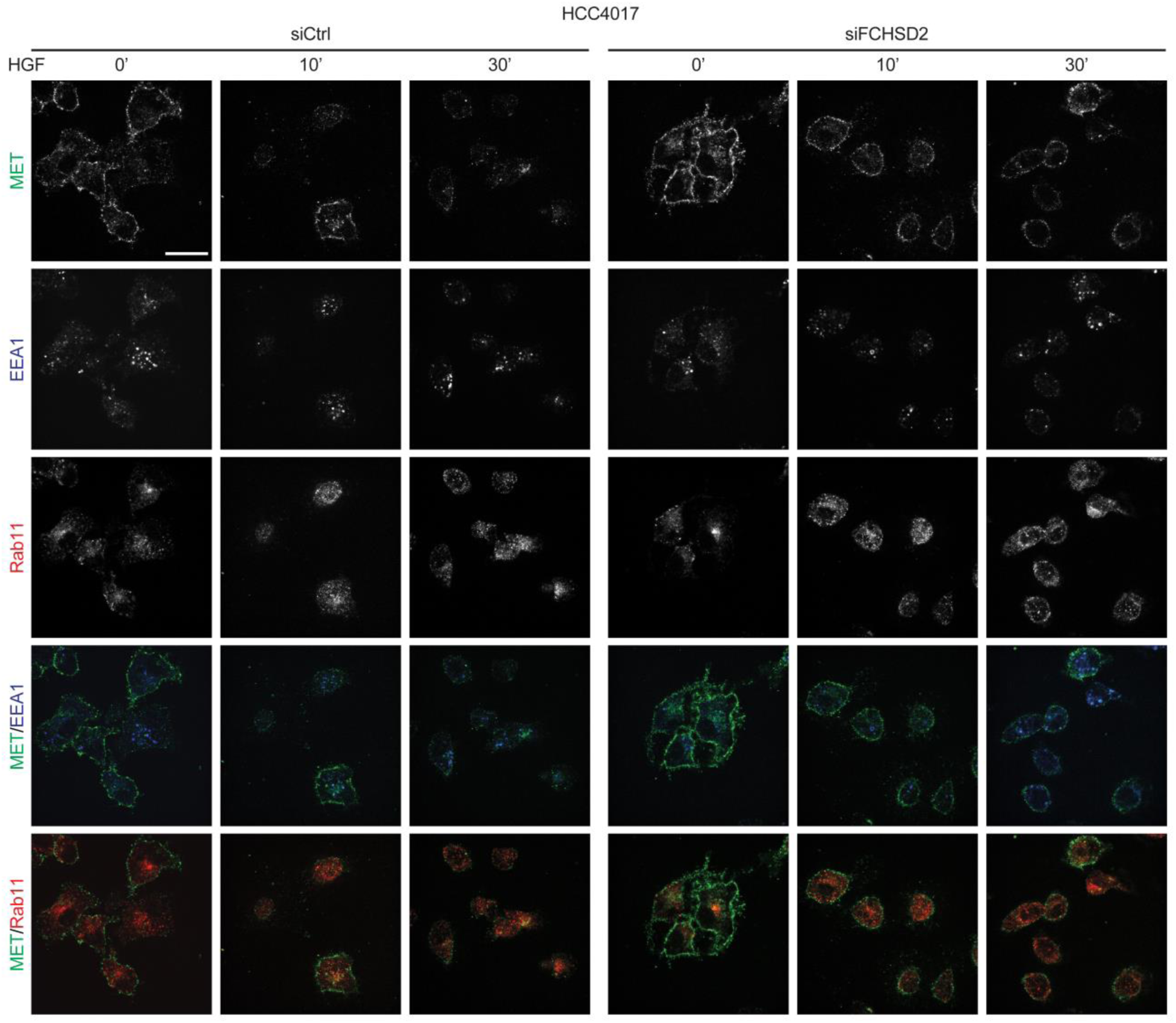
Representative confocal images of MET, EEA1 and Rab11 immunofluorescence staining in control or FCHSD2 siRNA-treated HCC4017 cells. Cells were incubated with 1 μg/ml HGF for 30 min at 4°C, washed, and reincubated at 37°C for the indicated times. Scale bar, 25 μm. Quantified results are shown in Fig. 2A.

**EV Figure 4.**
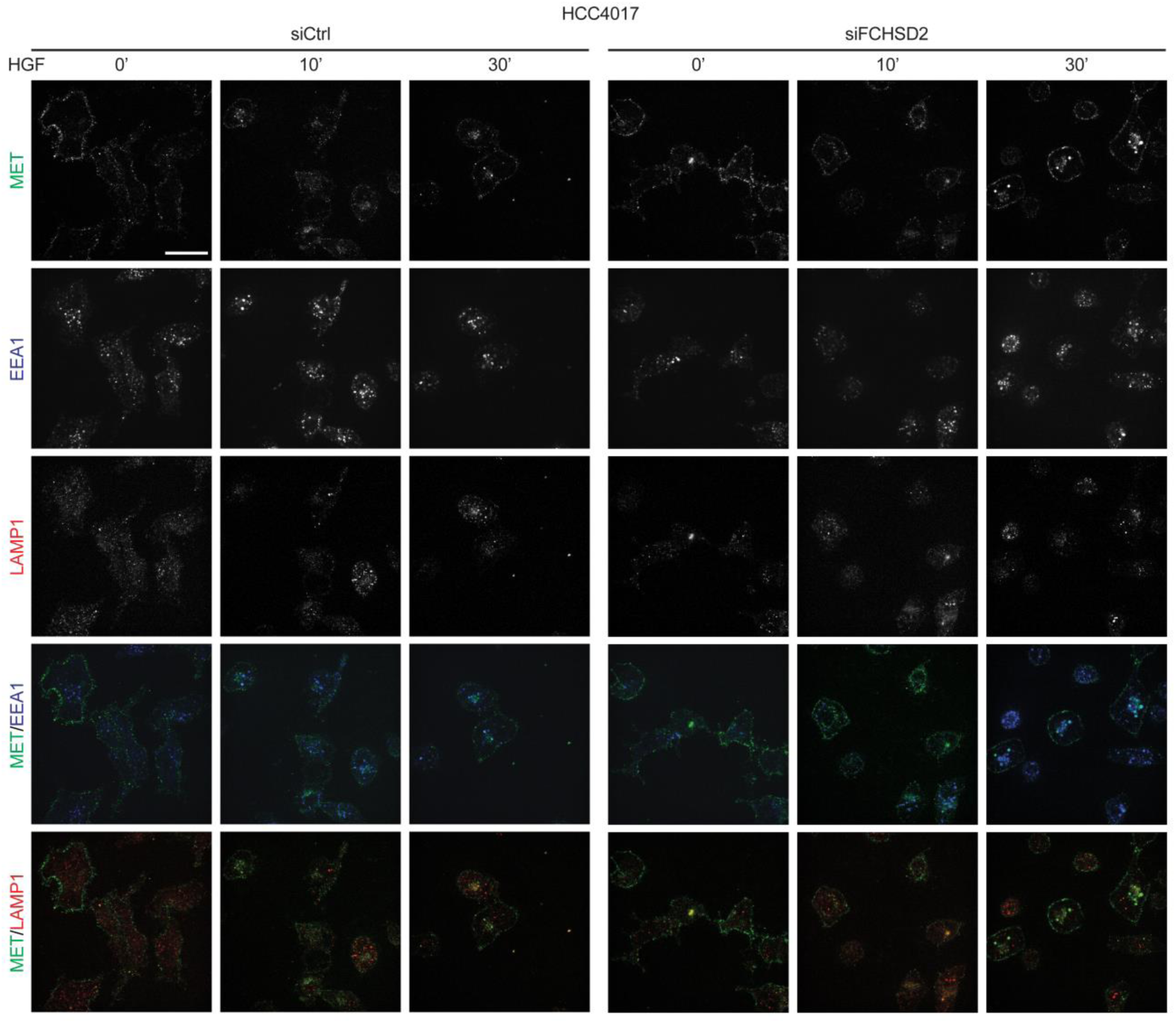
Representative confocal images of MET, EEA1 and LAMP1 immunofluorescence staining in control or FCHSD2 siRNA-treated HCC4017 cells. Cells were incubated with 1 μg/ml HGF for 30 min at 4°C, washed, and re-incubated at 37°C for the indicated times. Scale bar, 25 μm. Quantified results are shown in Fig. 2A.

**EV Figure 5.**
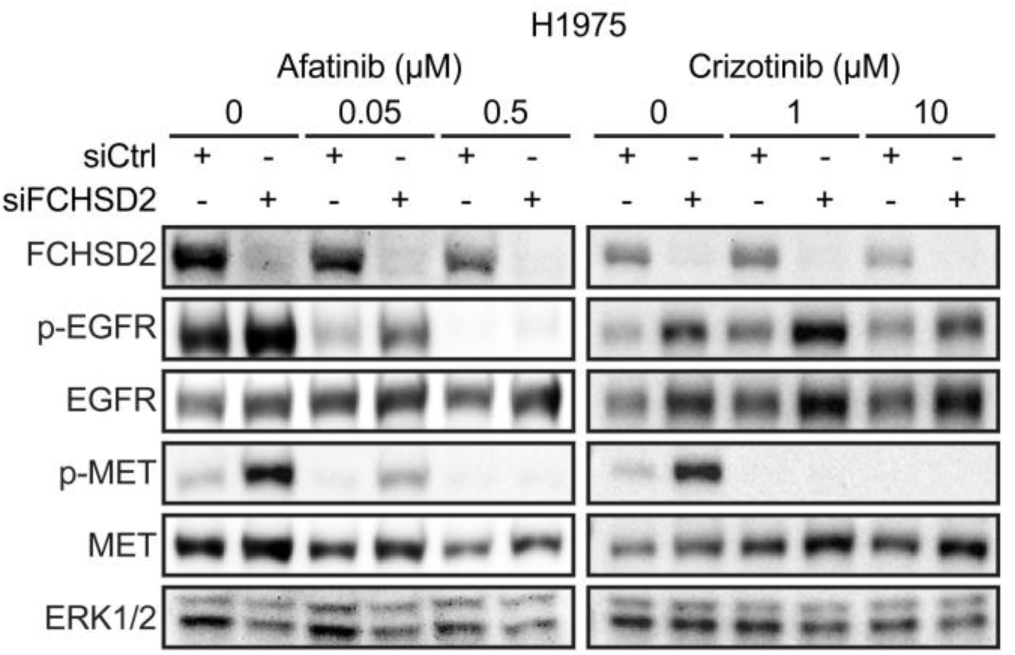
FCHSD2 depletion-induced upregulation of the RTKs is independent of their activities. H1975 control or FCHSD2 siRNA-treated cells were incubated with EGFR inhibitor (Afatinib) or MET inhibitor (Crizotinib) as indicated concentration for 24 h.

